## Supplemental Figures for "Resveratrol And Pterostilbene Potently Inhibit SARS-CoV-2 Replication In Vitro"

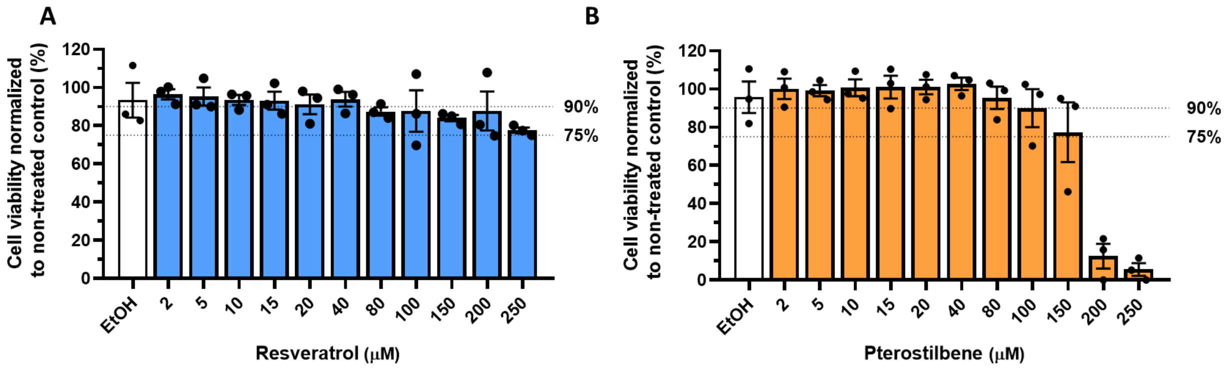


**Supplementary figure 1. Cellular cytotoxicity of resveratrol and pterostilbene in Vero E6 cells**. Cell viability of Vero E6 cells incubated with increasing concentrations of **(A)** resveratrol, **(B)** pterostilbene or **(A,B)** equivalent volumes of EtOH respective to the highest concentration of compound. Cell viability was assessed using an MTS assay kit. Cell viability is expressed as percentage compared to the non-treated (NT) control. Data are represented as mean ± SEM from three independent experiments.

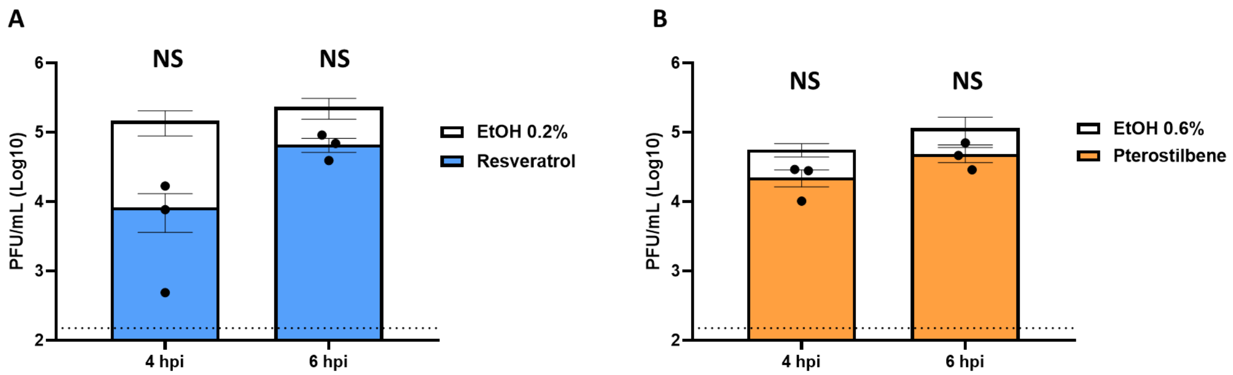


**Supplementary figure 2. Effect of resveratrol and pterostilbene when added late in the replication cycle of SARS-CoV-2.** Vero E6 cells were inoculated with SARS-CoV-2 at MOI 1 and treated with **(A)** 150 µM resveratrol, **(B)** 60 µM pterostilbene or **(A,B)** equivalent volumes of EtOH at 4 or 6 hpi. Virus production was determined at 8 hpi via plaque assay. Data is represented as mean ± SEM from three independent experiments. Dotted line indicates the threshold of detection. Student T test was used to evaluate statistical differences and a p value $\leq 0.05$ was considered significant with *$p \leq 0.05$, **$p \leq 0.01$ and ***$p \leq 0.001$ and NS as non-significant.

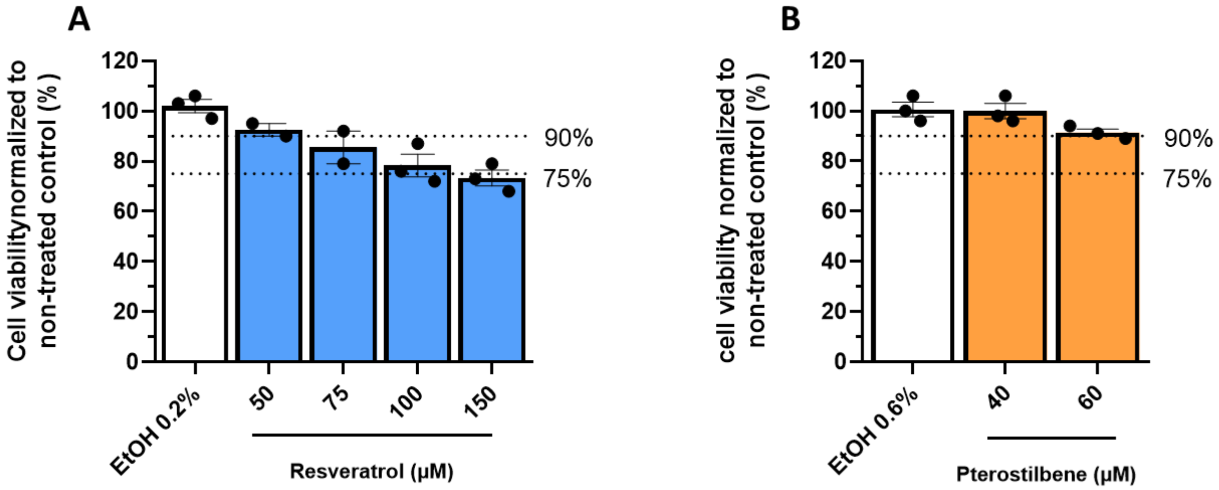


**Supplementary figure 3. Cellular cytotoxicity of resveratrol and pterostilbene in Calu-3 cells.** Calu-3 cells were incubated with increasing concentrations **(A)** resveratrol, **(B)** pterostilbene or **(A,B)** equivalent volumes of EtOH corresponding to the highest used concentration. Cell viability was assessed using an MTS assay. Cell viability is expressed as percentage compared to non-treated control. Data is represented as mean ± SEM from two or three independent experiments each performed in duplicates.

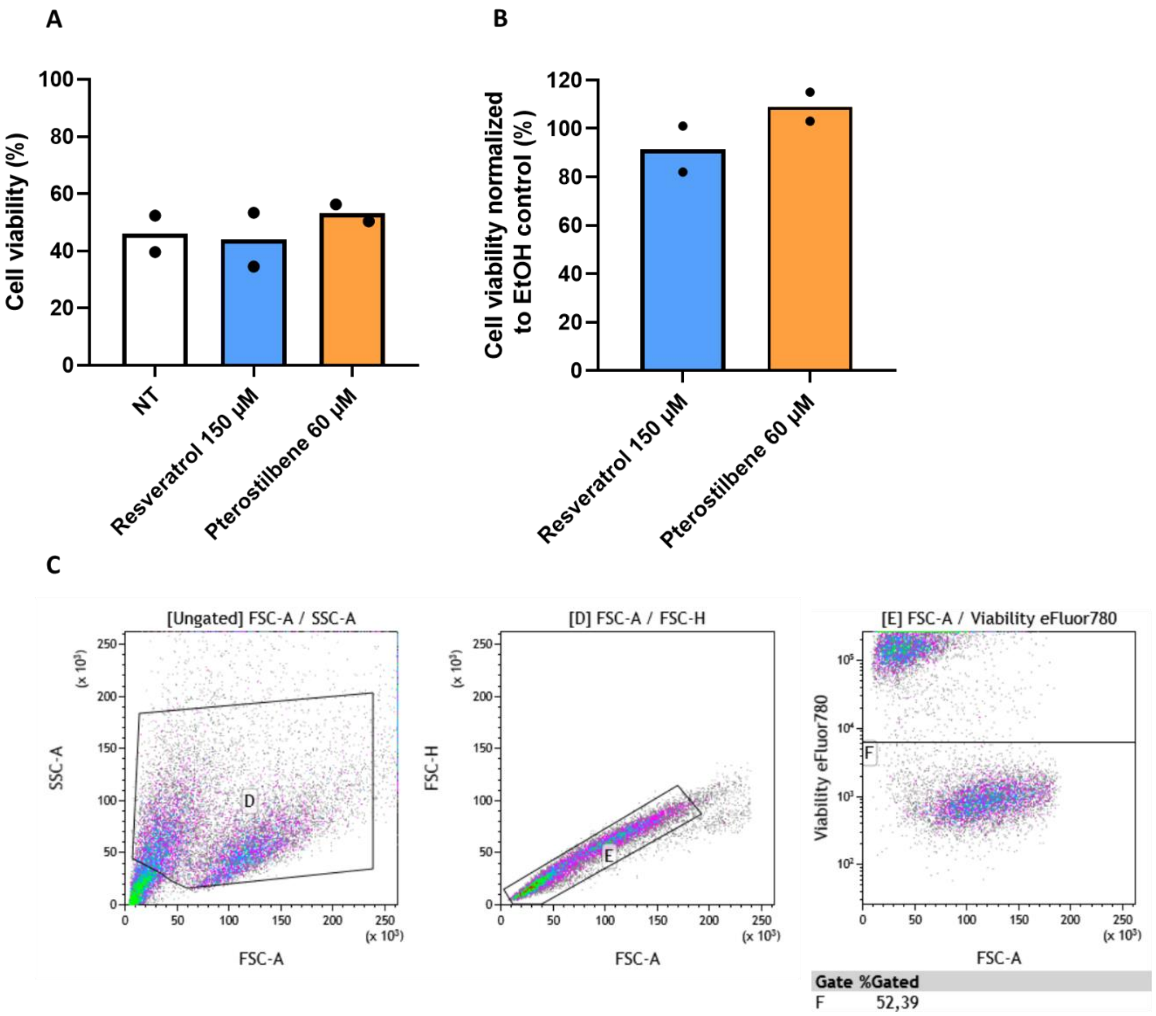


**Supplementary figure 4. Cellular cytotoxicity of resveratrol and pterostilbene in PBECs.** PBECs were incubated in absence (non-treated, NT) or presence of 150 µM resveratrol and 60 µM pterostilbene at basolateral side for 8 hr. Cell viability was assessed by flow cytometry using **(A)** life/death staining and **(B)** a LDH assay. **(C)** Gating strategy flow cytometry. Cell population was gated with gate D; doublets were excluded with gate E; live population was determined with gate F. Data represents mean ± SEM from two independent donors.
